# Chicdiff: a computational pipeline for detecting differential chromosomal interactions in Capture Hi-C data

**DOI:** 10.1101/526269

**Authors:** Jonathan Cairns, William R. Orchard, Valeriya Malysheva, Mikhail Spivakov

## Abstract

**Summary:** Capture Hi-C is a powerful approach for detecting chromosomal interactions involving, at least on one end, DNA regions of interest, such as gene promoters. We present Chicdiff, an R package for robust detection of differential interactions in Capture Hi-C data. Chicdiff enhances a state-of-the-art differential testing approach for count data with bespoke normalisation and multiple testing procedures that account for specific statistical properties of Capture Hi-C. We validate Chicdiff on published Promoter Capture Hi-C data in human Monocytes and CD4+ T cells, identifying multitudes of cell type-specific interactions, and confirming the overall positive association between promoter interactions and gene expression. Chicdiff is implemented as an R package that is publicly available at https://github.com/RegulatoryGenomicsGroup/chicdiff.

## 1. Introduction

Differential signal detection in sequencing data is one of the most common tasks in genomic analyses. Multiple tools have been developed for this purpose, many of which, including DESeq and EdgeR, are based on the negative binomial models for count data (Anders and Huber, 2010; Robinson *et al.*, 2010). Such tools are theoretically suitable for the analysis of most sequencing data types, including chromatin immunoprecipitation (ChIP-seq) and Hi-C, leading to the development of wrapper packages around DESeq and EdgeR that facilitate differential analyses for such data (Ross-Innes *et al.*, 2012; Lareau and Aryee, 2018). However, both of these algorithms have been developed with standard RNA sequencing data in mind, and may therefore not account for or benefit from the specific properties of data resulting from other assays.

Capture Hi-C (CHi-C) is a powerful experimental technique for detecting chromosomal interactions globally and at high resolution (Schoenfelder *et al.*, 2015). In CHi-C, the genome-wide pulldown of pairs of interacting genomic fragments by Hi-C is followed by sequence capture to selectively enrich Hi-C material for interactions involving (at least on one end) fragments of interest, termed ‘baits’. Differential analyses of CHi-C data are challenging due to sample normalisation issues, sparsity and uneven signal-to-noise ratios across interaction distances and different capture baits, which are not accounted for by standard differential analysis algorithms.

We have previously reported Chicago, a statistical pipeline for robust detection of significant interactions in Capture Hi-C data from a single condition (Cairns *et al.*, 2016). Here, we present Chicdiff, an R package for differential Capture Hi-C data analysis. Chicdiff combines moderated differential testing for count data implemented in DESeq2 (Love *et al.*, 2014) with CHi-C-specific procedures for signal normalisation informed by Chicago, and p-value weighting. Jointly, procedures implemented in Chicdiff enable a robust and sensitive detection of differential interactions in CHi-C data.

## 2. Approach

A schematic of the overall analysis approach is presented in Figure S1. The following sections and Supplementary Note describe specific steps in more detail.

### 2.1. Feature selection

CHi-C data are often sparse, particularly at large interaction distances, limiting the power of differential signal detection at single-fragment resolution even at significantly interacting regions. In part, this problem can be mitigated based on the fact CHi-C signals commonly spread to adjacent fragments (Eijsbouts *et al.*, 2018), most likely owing to the tethering of these fragments into the vicinity of the baits by nearby specific interactions. Therefore, to increase power, Chicdiff pools reads across several fragments (by default, five in each direction) surrounding each interacting fragment of interest for each bait. A functionality is provided to prioritise fragment-level interactions within each detected differentially interacting region post-hoc (see Supplementary Note).

### 2.2. Data normalisation and significance testing

Typically in differential count analyses, a single normalisation (scaling) factor is estimated per sample to account for differences in library size. However, we found that in CHi-C data, normalisation can be further improved by taking into account the differences in the background levels for specific pairs of fragments between samples. In CHi-C, unlike in many other data types such as RNA-seq, it is possible to obtain such background estimates from the data, and procedures for this are implemented in the Chicago package. Chicdiff combines scaling factors based on these background estimates with sample-level scaling factors in a manner that minimises the total dispersion of read counts across replicates and conditions at each interaction.

The count and scaling matrices generated as described above are provided as input for the DESeq2 package, which tests each interaction for differences between conditions using a negative binomial model with moderated dispersion estimation.

### 2.3. Weighted multiple testing treatment

As with other Hi-C-derived data types, signal-to-noise ratios and effect sizes in CHi-C data vary highly with interaction distance. This makes a strong case for non-uniform multiple testing correction, such that p-values for differential tests on longer-distance interactions are corrected more stringently compared with those on short-distance interactions. To do this, Chicdiff uses the Independent Hypothesis Weighting (IHW) method (Ignatiadis *et al.*, 2016) to learn p-value weights based on interaction distance in a manner that maximises the number of rejected null hypotheses. However, training IHW weights on the test regions is not appropriate since their p-values are often not uniform under the null due to selection bias, which violates IHW’s core assumption. Therefore, instead we learn weights on a separate ‘weight training set’ of fragment pairs randomly drawn from the full interaction count data for each sample (i.e., not limited to CHiCAGO-detected significant interactions), thus avoiding selection bias. The distance-dependent weights learned this way are applied to the p-values in the test set, and the resulting weighted p-values are reported to the user.

## 3. Use example

We applied Chicdiff to detect interactions specific to naive CD4+ T cells versus monocytes based on promoter CHi-C data from (Javierre *et al.*, 2016). This resulted in 208,232 detected differential interacting regions (weighted adjusted p-value < 0.05; see Table S1 for further summary statistics). An example of differential interactions is shown in Figure 1, and a heatmap of a subset of differential and non-differential interactions is shown in Figure S2. As expected, many genes whose promoters engaged in differential interactions showed consistent differences in expression between the two cell types (Figure S3).

**Figure 1.**
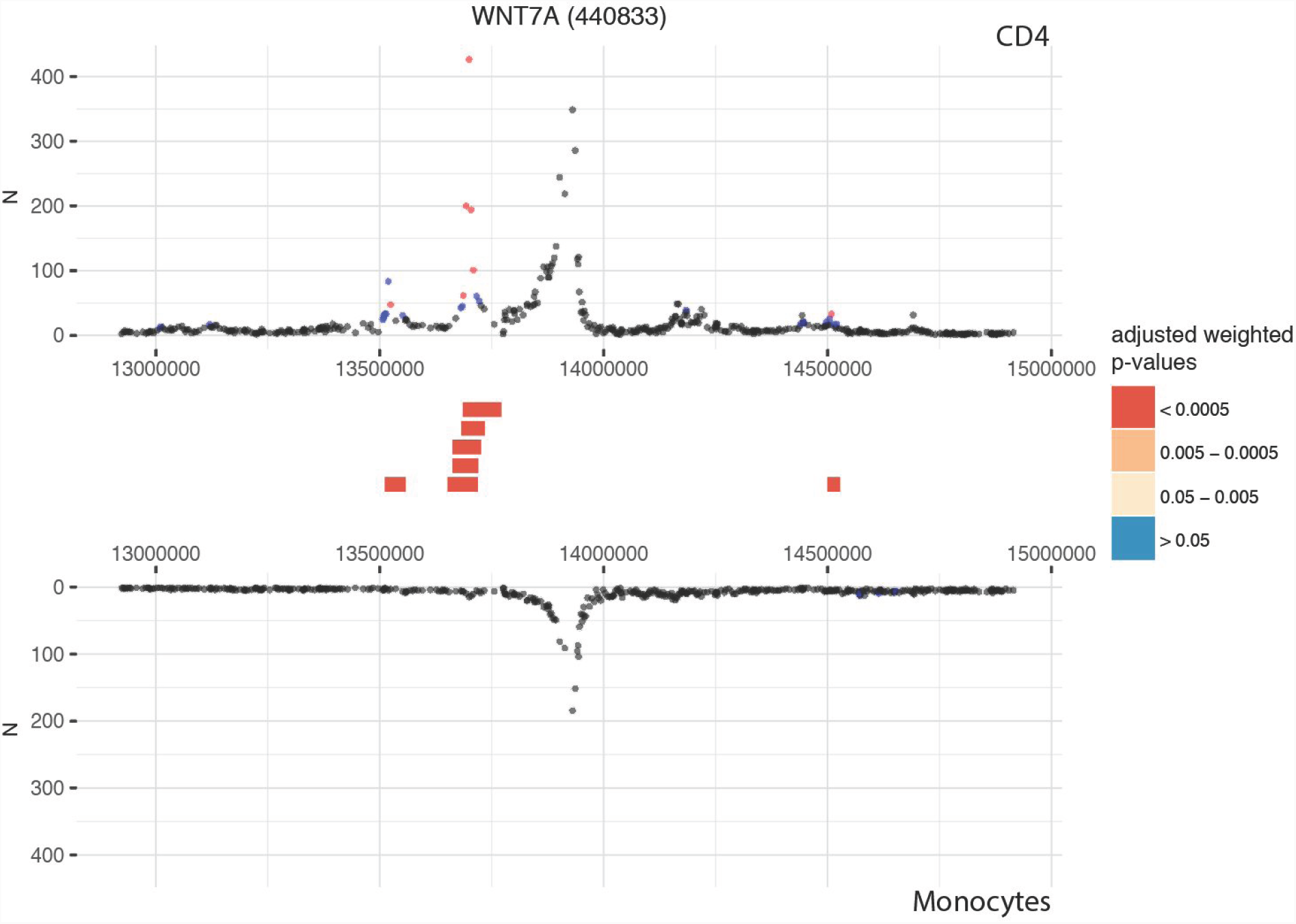
Example of differential interactions detected by Chicdiff. Profiles of Promoter Capture Hi-C interaction counts detected for *WNT7A* promoter in naive CD4+ T cells (top) and monocytes (bottom) (data from (Javierre *et al.*, 2016)). Mean counts across four and three replicates for each cell type, respectively, are shown along the Y axis, and interactions beyond 1Mb each way are cropped out. Significant interactions detected for each condition separately by Chicago are colour-coded (blue: 3<score<=5; red: score>5). Significant differentially interacting regions detected by Chicdiff (adjusted weighted p-value < 5e-4) are depicted as red blocks between the respective interaction profiles. The number in brackets in the plot title refers to the ID of the corresponding baited restriction fragment (440833).

Figures S4-S9 validate the Chicdiff approach by comparing the differential interaction calls obtained with and without pooling across multiple fragments, with Chicdiff versus standard DESeq2 normalisation, and with and without p-value weighting, with respect to the expression of associated genes and other parameters.

## 4. Conclusions

Capture Hi-C is a versatile experimental technique for detecting chromosomal interactions that involve, at least on one end, fragments of interest, such as gene promoters. Chicdiff extends and complements the Chicago statistical pipeline to provide a statistical framework for the detection of differential interactions between cell types and conditions in Capture Hi-C data. We expect Chicdiff to be widely used by the gene regulation and chromosome conformation communities.

## Acknowledgements

We thank Chris Wallace for helpful discussions and Michiel Thiecke for testing the developmental version of Chicdiff.

