## Supplementary note, Figures S1-10 and Table S1 for "Chicdiff: a computational pipeline for detecting differential chromosomal interactions in Capture Hi-C data"

Supplementary Information

Jonathan Cairns*, William R. Orchard*, Valeriya Malysheva*, and Mikhail Spivakov#

This document contains Supplementary Note, Figures S1-S10 and Table S1

### Supplementary Note

#### Extended description of Chicdiff methodology

***Input data.*** Chicdiff takes as input the output files produced by the Chicago pipeline that contain interaction-level estimates of the expected read count. Since Chicago uses several filtering steps, such as removing data for sparse baits, Chicdiff additionally expects a table of unfiltered read counts per fragment pair (.chinput files that are also used as input to the Chicago package). Finally, Chicdiff requires a list of interactions of interest, such as, for example, those with Chicago scores above a predefined cutoff (typically 5) in at least one replicate.

***Definition of the region universe.*** The first step is to define the features at the other end (OE) of each interaction of interest (the "region universe") to be used in the analysis. For most OEs, the regions are produced by extending the OE by a predefined number of fragments (by default, 5) in each direction. Overlapping regions generated from adjacent interactions of interest are allowed. Fragment extension is terminated if it hits the bait fragment or the end of the chromosome.

***Generation of the scaling matrix.*** Sample-level normalisation factors $s_k$ are generated by DESeq2 as follows:

$$s_k = \text{median } X_{ik}/X_i^R,$$

where $X_{ik}$ are read counts for a given interaction $i$ and sample $k$, and $X_i^R$ are those for the same interaction in the virtual "reference sample": $X_i^R = (\prod_k X_{ik})^{1/m}$, where $m$ is the total number of samples across all conditions.

Additionally, Chicdiff computes interaction-level normalisation factors $u_{ik}$. These are based on the estimates for the expected counts generated by the Brownian motion ($B_{ik}$) and technical noise ($T_{ik}$) for the corresponding bait-region interaction by the Chicago analysis of individual replicates:

$$u_{ik}^{raw} = 1/(B_{ik}+T_{ik}),$$

$$u_{ik} = u_{ik}^{raw} / (\prod_k u_{ik}^{raw})^{1/m}.$$

For each interaction $i$ and sample $k$, Chicdiff then seeks a mixture of sample-level and interaction-level normalisation factors ($s_k$ and $u_{ik}$):

$$S_{ik}^{raw} = s_k\theta + u_{ik}(1-\theta),$$
$$S_{ik} = S_{ik}^{raw} / (\prod_k S_{ik}^{raw})^{1/m}.$$

The mixing parameter $\theta$ is chosen such as to minimise the overall spread of normalised read counts at each interaction across all replicates and conditions. In practice, we minimise the sum of deviances of intercept-only DESeq2 regression models fitted at each interaction to the normalised read counts $X_{ik}^{norm} = X_{ik}/S_{ik}(\theta)$. For speed, we iterate over a discrete range of $\theta$, rather than minimising $\theta$ formally; minimising median deviance instead of the sum of deviances led to highly similar results (data not shown). For interactions, for which $B$ and $T$ are not available in at least one replicate and condition, because data for them were filtered out by Chicago ($i \in I_{filt}$), sample-level scaling factors $s_k$ are used instead (in other words, we override $\theta$ to 1 for all $i \in I_{filt}$).

This approach can be seen as similar in spirit (although not identical) to a shrinkage of interaction-level scaling factor estimates towards a “global” estimator, or simply as combining two estimators of the same parameter in the spirit of model or forecast averaging (Lavancier and Rochet 2016).

***Differential testing.*** Matrices of interaction-level counts and scaling factors are submitted to DESeq2’s moderated dispersion estimation procedure, followed by negative binomial regression using experimental condition as the explanatory variable. Wald test p-values are submitted to the weighted multiple testing procedure as described below.

***Weighted multiple testing procedure.*** Since the statistical power of the analysis (and likely also the true positive rate) strongly depends on distance, we weight p-values obtained in differential testing using the log-transformed interaction distance as the covariate, using an approach implemented in the R package IHW.

IHW expects the p-values to be uniform under the null; however in practice, this is often not the case for the interactions selected for Chicdiff testing due to selection bias. Therefore, we estimate weights on a ‘weights training control set’ of interactions that are randomly sampled from the full Capture Hi-C dataset in a manner that ensures a sufficient representation of interactions across the whole range of distances, in terms of the number of spanned restriction fragments (based on the normal distribution with a mean of zero and a standard deviation equal to one third of the maximum distance observed for interactions in the test set). Interactions from the weights training set are pooled into regions, normalised and assigned p-values in the same way as those in the test set. The only exception is that the scaling factor mixing parameter $\theta$ is not re-estimated, and instead the value of $\theta$ optimised on the test set is used in normalisation.

P-values for interactions in the test set are adjusted based on the distance-dependent weights learned this way, followed by Benjamini-Hochberg (BH) multiple testing correction.

***Prioritisation of fragment-level interactions.*** It may be desirable to obtain information at the level of individual fragments, as opposed to pooled regions detected as differential. To this end, Chicdiff first combines the p-values for all regions corresponding to a given fragment-level interaction. By default, it takes the minimum BH-adjusted weighted p-value over all regions, but incorporates the option to combine p-values formally using the Harmonic Mean P-value approach for dependent tests (Wilson 2019), which is slightly more conservative (Figure S10). To prioritise the potential 'driver' interactions within each region, we provide the functionality to filter them by the differences in mean asinh-transformed Chicago interaction scores between conditions. Asinh transformation is performed to emphasise differences within the low range of scores (such as those between scores of 0 and 6, given the typically used Chicago score cutoff of score>=5), as such differences are more interpretable compared with those in the high range (such as those between scores of 20 and 25).

***Visualisation of the results.*** Chicdiff provides a plotting function to visualise the detected differentially interacting regions. Interaction-level mean read counts for a given bait are plotted for the two conditions as mirror images. Differentially interacting regions are depicted in the space between the two profiles, colour-coded by adjusted weighted p-value. This plotting function was used to generate Figure 1.

**Chicdiff implementation and resource requirements**

Chicdiff is implemented as an R package using Bioconductor R packages Chicago, DESeq2 and IHW as key dependencies, requiring R version >=3.4.3. Resource requirements differ depending on the number of replicates and the read coverage of the samples. The high-coverage example dataset in this paper (3 replicates of monocytes and 4 replicates of naive CD4 T cells, with each replicate sequenced across 3 lanes on an Illumina HiSeq2500 machine) uses ~40 Gb RAM and takes ~2 hrs to complete on a standard Linux compute node. For training purposes, we provide a subset of the data from the use case (chr19 only; two replicates for each condition) in the data package ChicdiffData, which uses ~0.6 Gb RAM and takes ~5 min to process with Chicdiff on a standard laptop machine.

**Data and code for the use example**

The datasets for the use example and the code used to run Chicdiff and generate figures have been deposited to https://osf.io/y9nb5/.

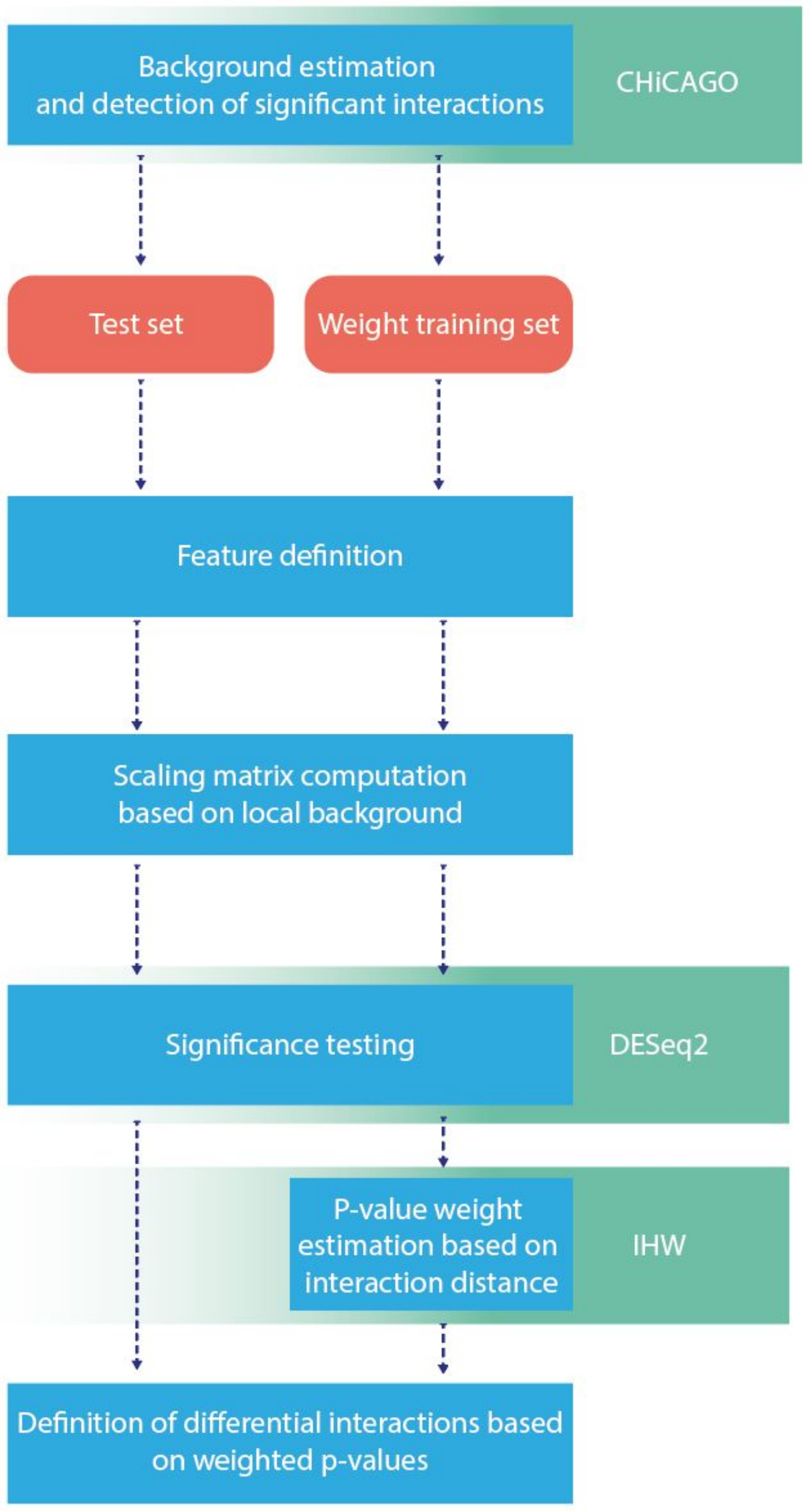


**Figure S1. Schematic of the Chicdiff analysis approach.** See main text and Supplementary Note for details.

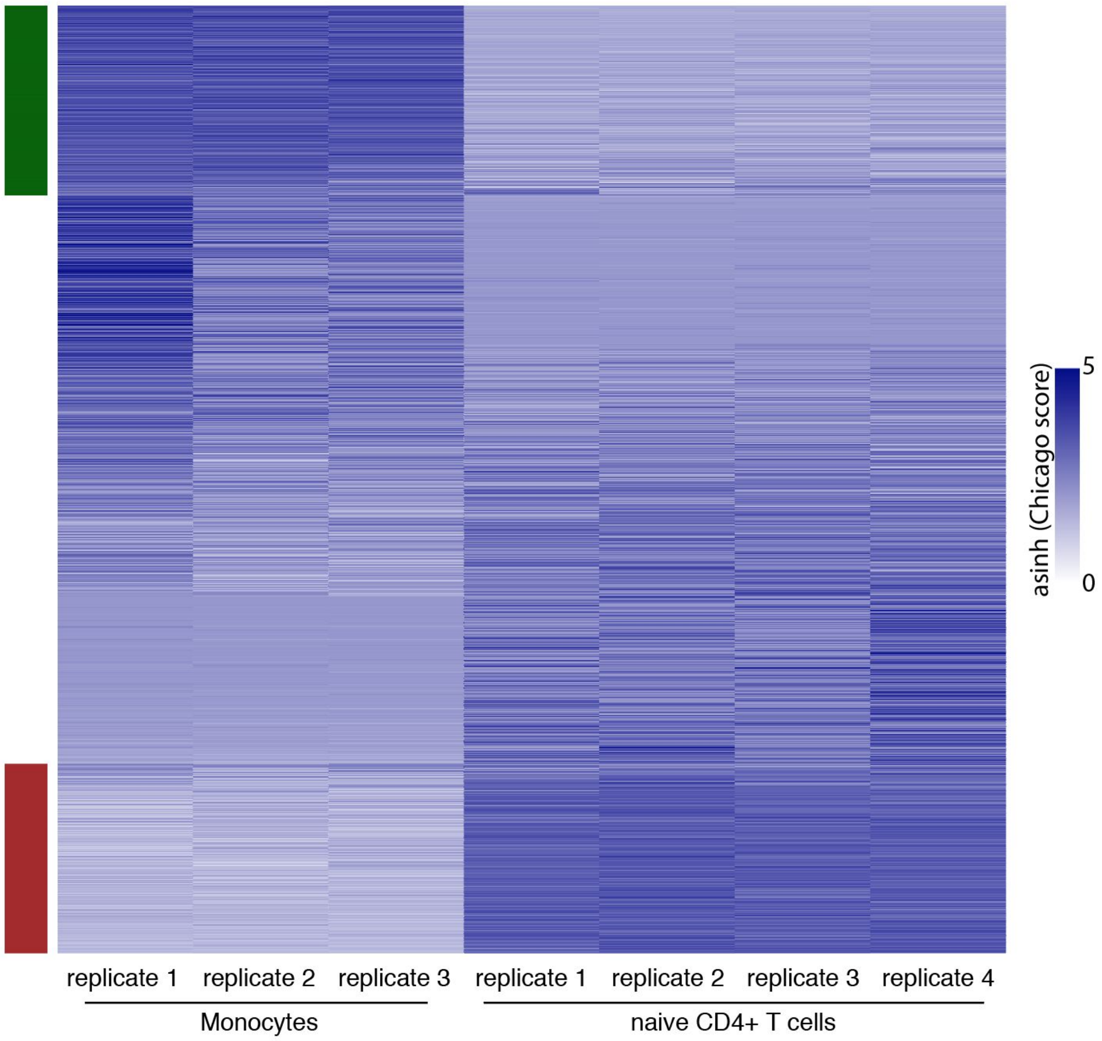


**Figure S2. Heatmap of a subset of differential and non-differential interactions.** Heatmap presenting a random subset of 1000 promoter interactions mapping within the detected differentially interacting regions (min [adjusted weighted p-value across a set of regions containing the fragment of interest] < 1e-3) between Monocytes and naive $CD4^+$ T cells, as well as 3000 interactions not detected as differential (min [adjusted weighted p-value across a set of regions containing the fragment of interest] > 0.5). Asinh-transformed Chicago scores are colour-coded on a scale from white to dark blue. Strips to the left of the heatmap demarcate interactions mapping within differentially interacting regions, with a higher Chicago score observed in monocytes (dark green) and naive $CD4^+$ T cells (purple), respectively.

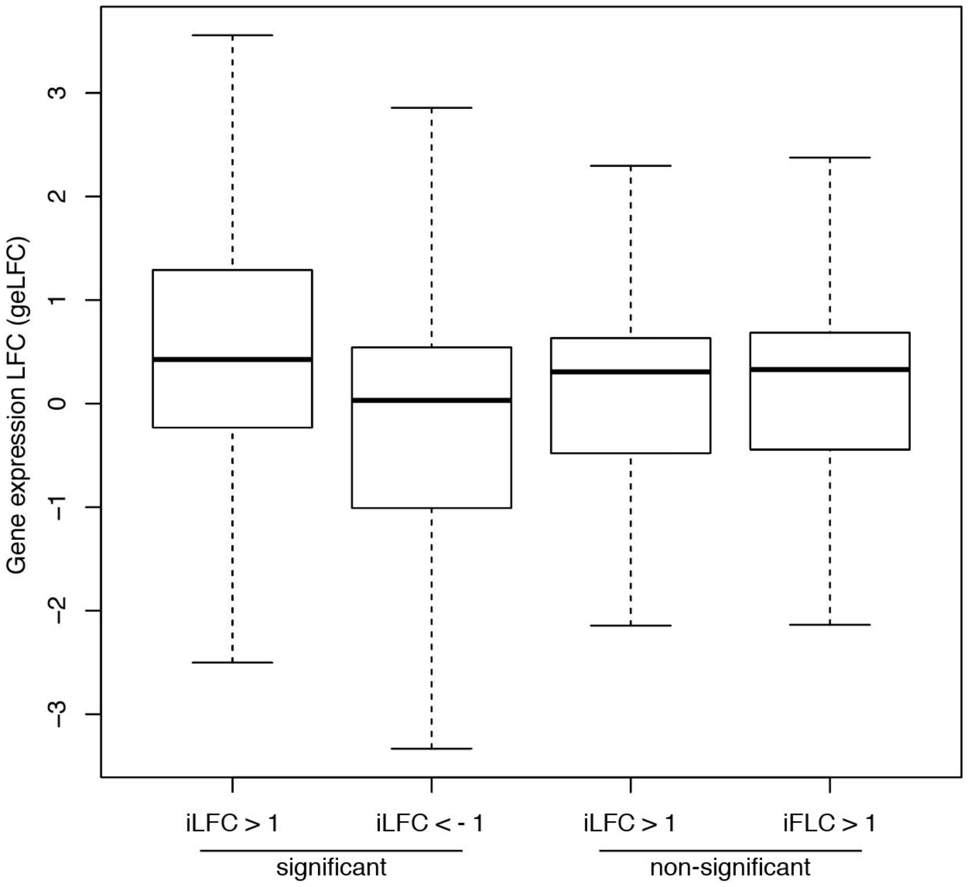


**Figure S3. Differential promoter interactions associate with differential gene expression of the respective genes.** Boxplots showing changes in the expression levels (on the log scale, geLFC) for genes associated with significant vs non-significant differential interactions between promoters and pooled regions detected with Chicdiff (adjusted weighted p-values below 1e-5 and above 0.05, respectively), and having a $\log_2$ fold change in normalised interaction read counts (iLFC) of at least 1 in either direction. The distributions of geLFCs of genes associated with significant differential interactions were shifted in the same direction as the respective iLFCs (Wilcoxon test p-value=1e-113; for genes with more than one associated differential interaction, one interaction was chosen for testing to avoid inflating the significance due to correlated observations). In contrast, the distributions of geLFCs for genes associated with non-significantly different interactions showed only a very minor shift that was in the opposite direction with respect to the respective iLFCs. Outliers were omitted for clarity. MMSEQ-estimated relative gene expression levels for monocytes and naive $CD4^+$ T cells were from the BLUEPRINT project as released in (Javierre et al. 2016).

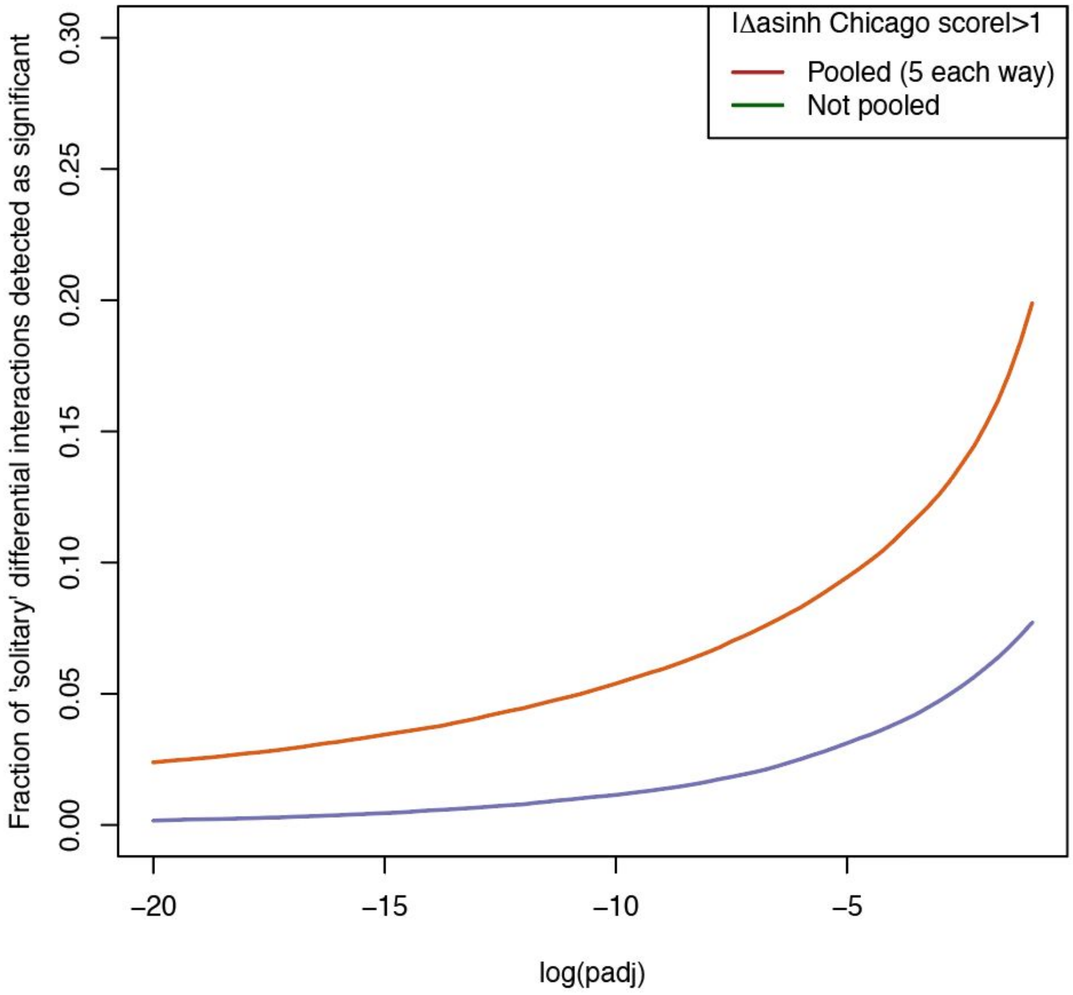


**Figure S4. Increase in sensitivity owing to pooling adjacent fragments into test regions.** Fraction of interactions with |△ mean asinh-transformed Chicago scores|>1 that were detected as differential over a range of $\log_{10}$-transformed adjusted weighted p-value thresholds (brown: pooling five adjacent fragments each way into test regions; dark green: no pooling). To minimise the effect of ‘passenger’ interactions within the pooled regions, we considered only ‘solitary’ interactions, whereby only a single interaction had the above difference in Chicago scores, and at most one was detected as differential without fragment pooling at a given log p-value threshold.

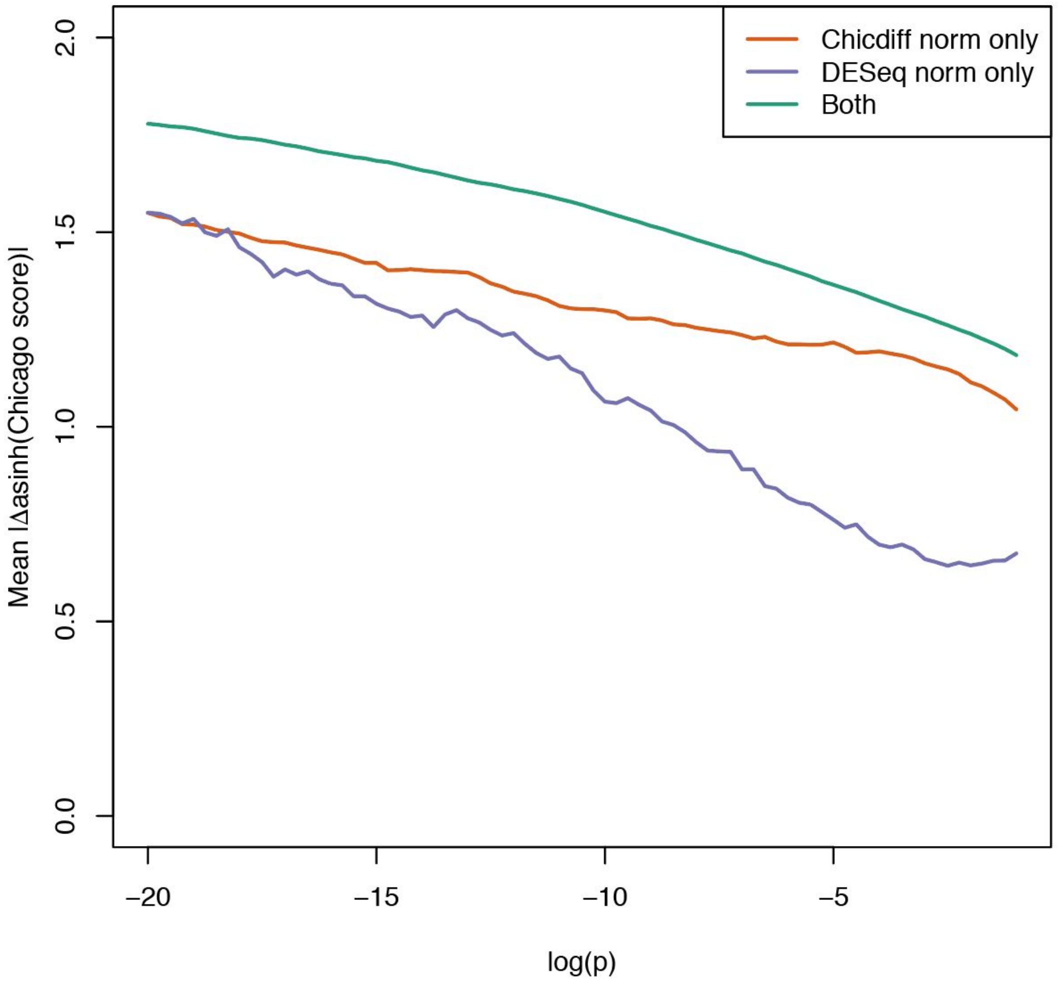


**Figure S5. Comparison of interactions detected using Chicdiff versus standard DESeq normalisation.** The means of |△ mean asinh Chicago scores between conditions| across interactions that exceed a given $\log_{10}$ p-value cutoff with Chicdiff normalisation only (brown; Chicdiff-estimated scaling factor mixing parameter $\theta$=0.75), DESeq normalisation only (blue; equivalent to $\theta$=1) or both approaches. These results suggest that while the strongest differential signals are generally detected irrespective of the normalisation approach, those detected with Chicdiff normalisation alone are likely to be stronger than those detected with DESeq alone. The figure is based on unweighted, unadjusted p-values to avoid confounding by multiple testing treatment (using weighted, adjusted p-values produces a highly similar result; not shown).

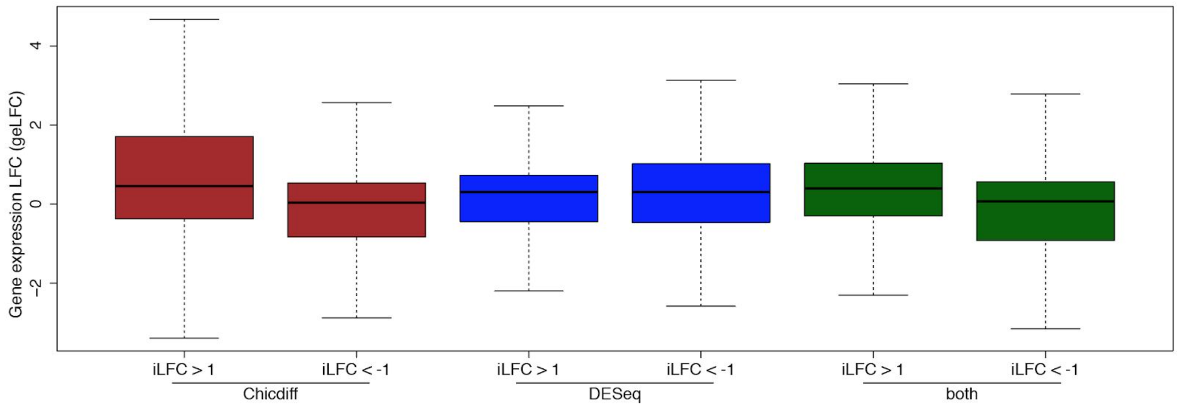


**Figure S6. Differences in the expression of genes associated with differential interactions detected with Chicdiff and standard DESeq normalisation.** Boxplots showing changes in the expression levels (on the log scale, geLFC) for genes associated with significant differential interactions between promoters and pooled regions detected with Chicdiff normalisation, DESeq normalisation or both approaches, respectively, (adjusted weighted p-value cutoff 1e-3), and having a $\log_2$ fold change in normalised interaction read counts (iLFC) of at least 1 in either direction. The distributions of geLFCs for genes associated with significant differential interactions detected with either Chicdiff normalisation alone or both normalisation approaches, the geLFC distributions for the associated genes were shifted in the same direction as iLFCs (Wilcoxon test p-values of 5e-16 and 2e-74, respectively; for genes with more than a single associated differential interaction, one interaction was chosen for testing to avoid inflating the significance due to correlated observations). In contrast, the distributions of geLFCs for genes associated with differential interactions detected with DESeq normalisation alone were generally similar irrespective of the direction of the respective iLFCs (Wilcoxon test p-value=0.92). Outliers were omitted for clarity. MMSEQ-estimated relative gene expression levels for monocytes and naive $CD4^+$ T cells were from the BLUEPRINT project as released in (Javierre et al. 2016).

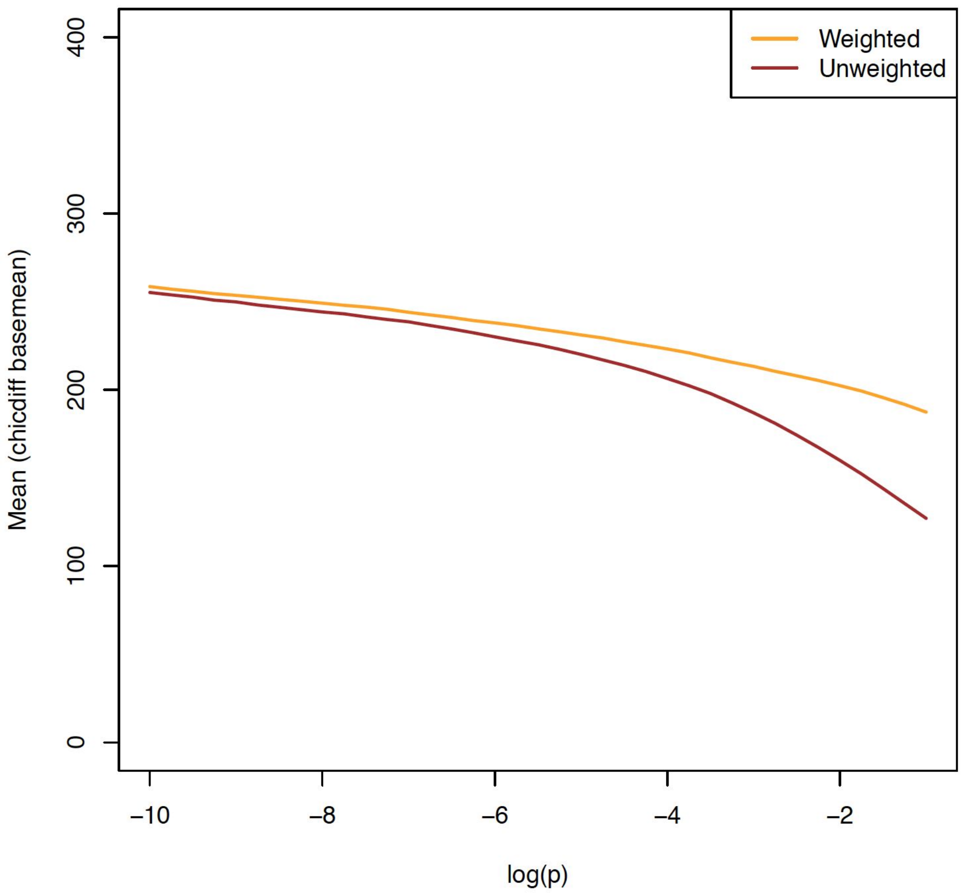


**Figure S7. P-value weighting prioritises differential interactions associated with a higher mean read count.** The mean base mean levels of interactions detected over a range of thresholds on either unweighted (orange) or weighted (brown) $\log_{10}$ p-values. Interactions with $\log_{10}$-p-values below –10 are not shown, as differences between the mean counts for weighted and unweighted p-values in this range are nearly indistinguishable, as can be extrapolated from the observed trend.

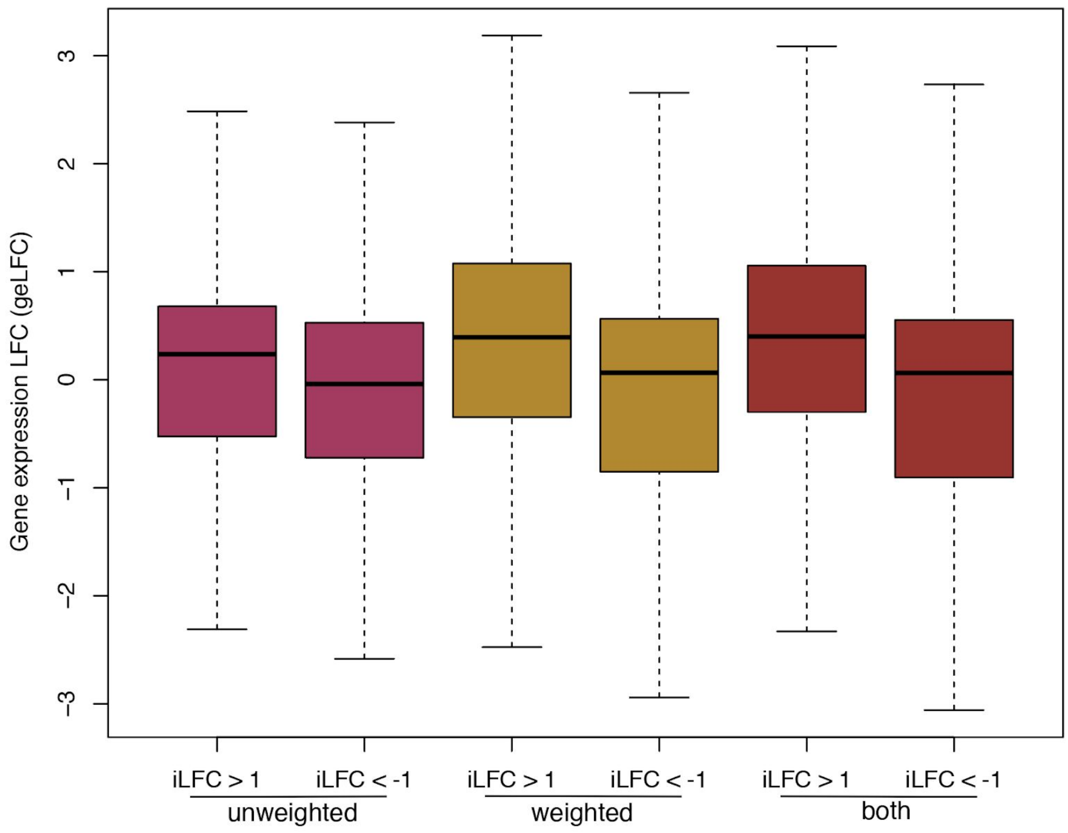


**Figure S8. Differential promoter interactions detected with p-value weighting associate with genes showing larger differences in expression.** Boxplots showing changes in the expression levels (on the log scale, geLFC) for genes associated with significant differential interactions between promoters and pooled regions detected with or without p-value weighting, or with both approaches (adjusted p-value<1e-3), and having a $\log_2$ fold change in normalised interaction read counts (iLFC) of at least 1 in either direction. The distributions of geLFCs for genes associated with significant differential interactions were shifted in the same direction as the respective iLFCs. This shift was markedly stronger when interactions were detected using weighted p-values (or both approaches) compared with those detected without weighting. Outliers were omitted for clarity. MMSEQ-estimated relative gene expression levels for monocytes and naive $CD4^+$ T cells were from the BLUEPRINT project as released in (Javierre et al. 2016).

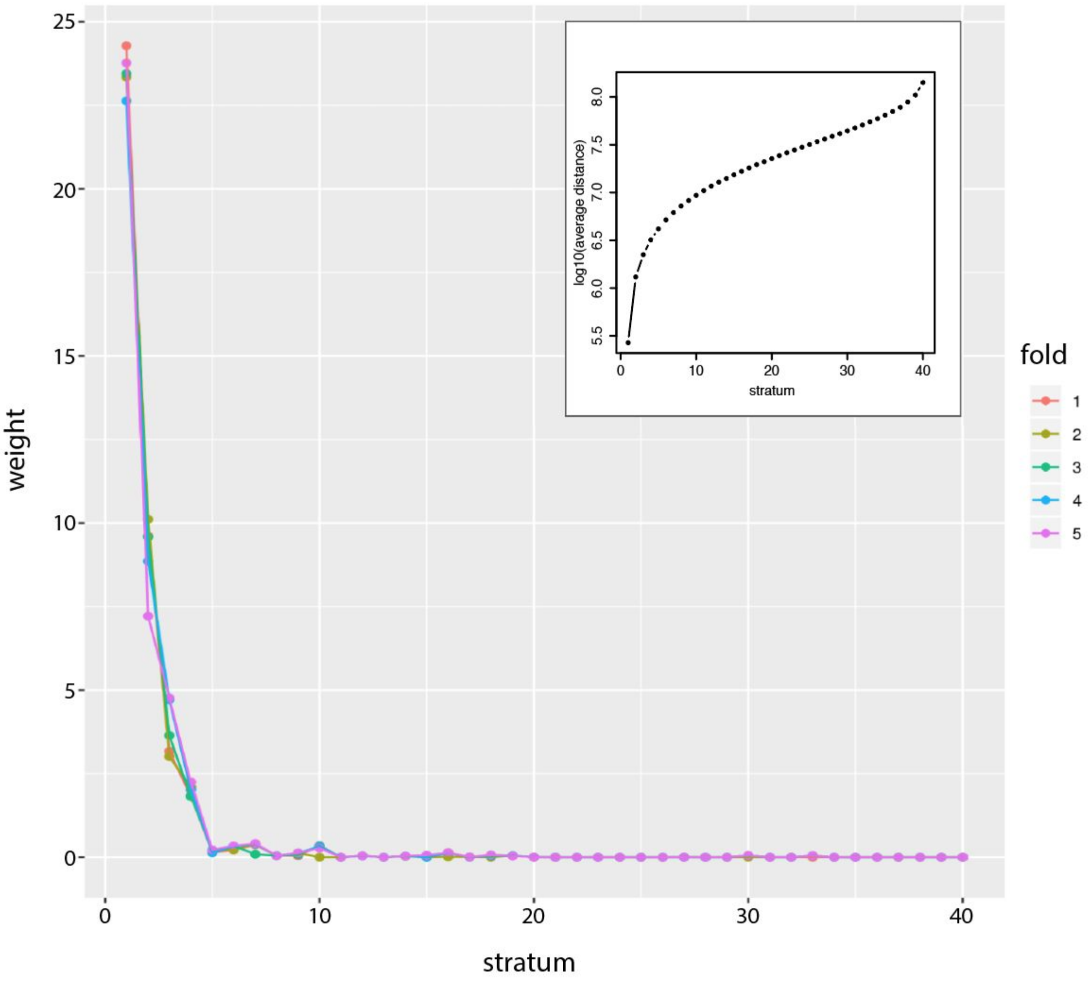


**Figure S9. P-value weights identified by IHW using interaction distance as a covariate.** The p-value weights (Y axis) learned by the IHW algorithm for each stratum of log-interaction distance over five cross-validation folds. The main plot has been generated by the IHW package invoked by Chicdiff. The inset shows the average interaction distance ($\log_{10}$-bps) for each stratum.

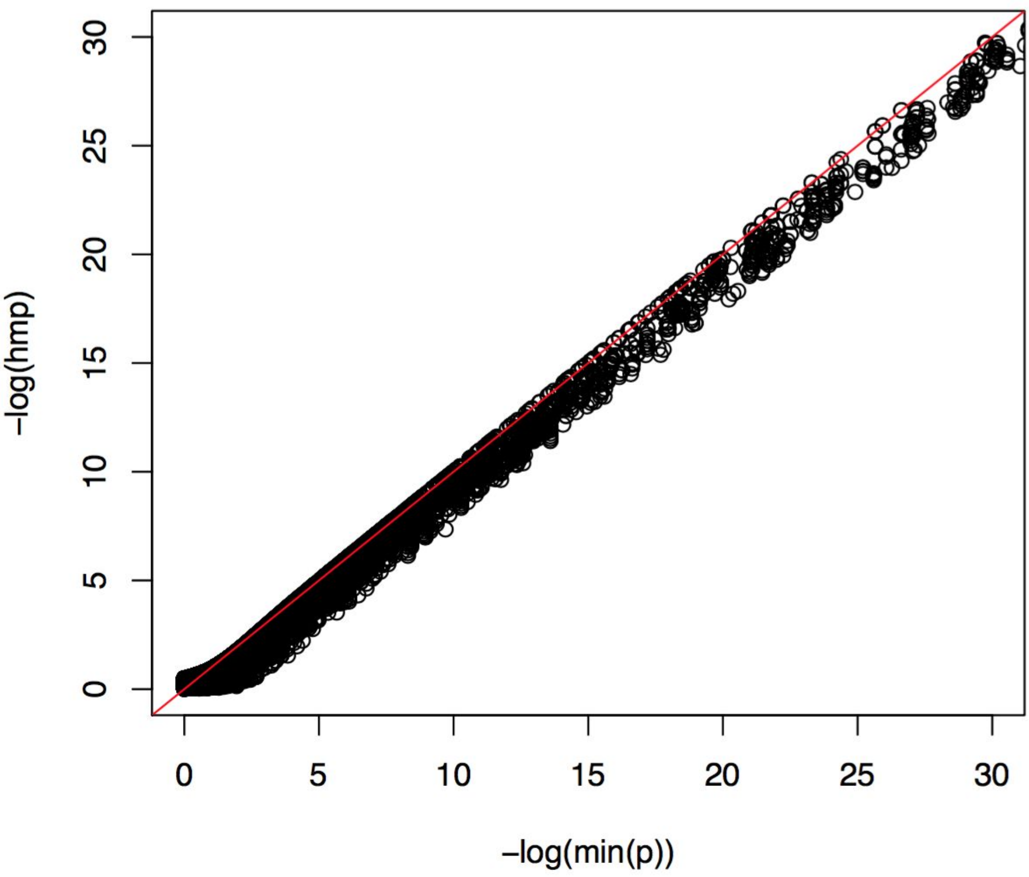


**Figure S10. Comparison of two strategies for inferring fragment-level p-values.** A scatterplot of fragment-level p-values for interactions on chr19 combined from the pooled region-level weighted, adjusted p-values using either the minimum p-value (X axis) or the harmonic mean approach (Wilson 2019) (hmp, Y axis). The y=x line is shown in red.

**Table S1. Summary statistics for the use example**

| Samples | Naive CD4 T cells (4 replicates) vs monocytes (3 replicates) | | | |
|---|---|---|---|---|
| Interactions tested | 831,052<br>(Chicago score > 5 in at least one replicate and cell type) | | | |
| Fragment pooling | 5 each way | None | 5 each way | None |
| Normalisation | Chicdiff<br>(optimised θ=0.75) | | Standard DESeq2<br>(equivalent to θ=1) | |
| No. differential regions (unweighted padj<0.05) | 262,398 | 81,064 | 230,294 | 74,164 |
| No. differential regions (weighted padj<0.05) | 208,187 | 71,257 | 184,583 | 69,476 |
| No. prioritised differential fragments (min weighted padj<0.05; $\lvert\Delta$ mean asinh-Chicago score$\rvert>1$) | 122,274 | 56,536 | 106,523 | 58,049 |
